# The Master Database of All Possible RNA Sequences and Its Integration with RNAcmap for RNA Homology Search

**DOI:** 10.1101/2023.02.01.526559

**Authors:** Ke Chen, Thomas Litfin, Jaswinder Singh, Jian Zhan, Yaoqi Zhou

**Author notes:** Co-first authors (Ke Chen, Thomas Litfin).

## Abstract

Recent success of AlphaFold2 in protein structure prediction relied heavily on co-evolutionary information derived from homologous protein sequences found in the huge, integrated database of protein sequences (Big Fantastic Database). In contrast, the existing nucleotide databases were not consolidated to facilitate wider and deeper homology search. Here, we built a comprehensive database by including the noncoding RNA sequences from RNAcentral, the transcriptome assembly and metagenome assembly from MG-RAST, the genomic sequences from Genome Warehouse (GWH), and the genomic sequences from MGnify, in addition to NCBI’s nucleotide database (nt) and its subsets. The resulting MARS database (Master database of All possible RNA sequences) is 20-fold larger than NCBI’s nt database or 60-fold larger than RNAcentral. The new dataset along with a new split-search strategy allows a substantial improvement in homology search over existing state-of-the-art techniques. It also yields more accurate and more sensitive multiple sequence alignments (MSA) than manually curated MSAs from Rfam for the majority of structured RNAs mapped to Rfam. The results indicate that MARS coupled with the fully automatic homology search tool RNAcmap will be useful for improved structural and functional inference of noncoding RNAs.

## INTRODUCTION

There are two major categories of RNAs: those coded for proteins (messenger RNA, mRNA) and those not (noncoding RNAs, ncRNA). The first ncRNA discovered was transfer-RNA (tRNA) in 1958[1]. Since then, new types of noncoding RNAs were constantly uncovered once every a few years[2]. These noncoding RNAs can have a length ranging from ~20 nucleotides in microRNAs (miRNA)[3] to >100kB for a long noncoding RNA (lncRNA) like AIR [4]. These RNAs can perform functions at the sequence level by simple complementary base-pairing in the case of miRNA[3], at the secondary structural level in the case of RNA switches [5], and at the tertiary structural level in the cases of tRNA, ribosomal RNA (rRNA), ribozymes, and riboswitches [6]. The number of distinct ncRNAs greatly exceeds that of distinct proteins. This is exemplified by the fact that our human genome dedicated more than 70% for RNA transcripts, compared to a tiny 1.5% coded for proteins [7]. These ncRNAs actively participate in essentially all biological processes and implicated in >1000 diseases[2,8]. Given increasingly importance for annotated and unannotated RNAs in biology (coding and noncoding), a comprehensive sequence database for all RNAs is necessary.

The most comprehensive database for ncRNAs is perhaps RNAcentral [9], which consolidates 56 Expert Databases and over 30 million sequences as of Jan 2022 (release 20). Another widely used sequence library is NCBI’s nucleotide database (nt) [10]. Unlike RNAcentral, NCBI’s nt database contains both RNA and DNA sequences. It combined the GenBank, European Nucleotide Archive (EMBL-EBI), and DNA Data Bank of Japan databases with a sequence count of 72.9 million as of Aug 2021. However, neither RNAcentral, nor NCBI’s nt database is complete for all possible RNA sequences as many specialized databases and depositories are not included.

Recently, AlphaFold2 achieved an incredible feat of accurate protein structure prediction for most predicted proteins in the 14^th^ biannual meeting of critically assessment of structure prediction techniques (CASP 14) [11]. This success was in part built on utilization of homologous sequences to extract evolution and co-evolution information, which contains implicitly the information on sidechain-sidechain distances and backbone/sidechain torsion angles. To secure as many homologous sequences as possible, they utilized the Big Fantastic Database (BFD) covering over 2 billion protein sequences from reference databases, metagenomes and metatranscriptomes.

Inspired by BFD, we build the Master database of All possible RNA Sequences (MARS). As in the nt database, we incorporated both RNA and DNA sequences, including genomic sequences. Genomic sequences were included because a large portion of genomic sequences are transcribed into coding and noncoding RNAs. Their inclusions allow us to account for all possible (or potential) RNAs.

To illustrate the usefulness of the MARS database, we compare the ability to obtain homologous sequences by using the fully automatic pipeline RNAcmap [12]. In this RNAcmap pipeline, a query sequence is first searched against a database by Blast-N [13], followed by a covariance-model-based search by Infernal [14]. Resulting multiple sequence alignment (MSA) was then evaluated by direct coupling analysis tools such as mfDCA [15]. Evolution and co-evolution information obtained from RNAcmap were found useful in improving RNA secondary structure and tertiary base-pair prediction in SPOT-RNA2 [16] as well as distance-contact map prediction in SPOT-RNA-2D [17]. In the latest update of RNAcmap (RNAcmap2) [18], an additional search by Infernal was performed on the multiple sequence alignment (MSA) produced by RNAcmap. A slightly expanded database was also utilized in RNAcmap2 by including environment samples (env nt), transcriptome shotgun assembly (tsa nt), and nucleotide sequences derived from the Patent Division of GenBank (pat nt) databases in addition to NCBI’s nucleotide (nt) database. The additional iteration as well as the database expansion were found effective in improving the quality of MSA obtained by examining the accuracy of base pairs extracted from the MSA using direct coupling analysis[18]. More recently, an rMSA pipeline was also proposed[19] and found useful in predicting RNA distance and orientation maps by deep learning [20]. It performed five iterative searches against Rfam [21], RNAcentral [9], the nt database [10] by using Blast-N[13], nhmmer[22] and Infernal [14].

Here, we established MARS database by incorporating several additional resources. They include the RNAcentral database [9], the transcriptome assembly and metagenome assembly hosted at the University of Chicago (MG-RAST) [23,24], the genomic sequences from Genome Warehouse (GWH)[25,26] and the genomic sequences from MGnify[27]. This database has more than 20 folds (or 60 folds) over the number of sequences in NCBI’s nt database (or the RNAcentral database). We illustrated the usefulness of MARS by employing a data splitting strategy coupled with the homology search tool RNAcmap2. The resulting RNAcmap3 increases 36 folds in the median number of effective homologous sequences and 2 folds in the F1-score for base pair prediction by direct coupling analysis over RNAcmap2 for no-hit RNAs (those RNAs lacking homologs according to RNAcmap1). RNAcmap3 also yields more accurate multiple sequence alignments (MSA) than rMSA and manually curated MSAs from Rfam for the majority of structured RNAs mapped to Rfam.

## DATA AND METHOD

### Data collection

The MARS database integrates all available nucleotide sequences, ranging from well-annotated individual nucleotide sequences to poorly understood metagenomics assemblies. Specifically, the data source of MARS includes NCBI’s nucleotide database (nt)[10], environmental samples (env_nt)[10], transcriptome shotgun assembly (tsa_nt)[10] and nucleotide sequences from the Patent Division of Genbank (patnt)[10], the noncoding RNA sequences from RNAcentral[9], the transcriptome assembly and metagenome assembly from MG-RAST[23,24], the genomic sequences from Genome Warehouse (GWH)[25,26] and the genomic sequences from MGnify[27].

The nt, env_nt, tsa_nt and patnt databases were downloaded from ftp://ftp.ncbi.nlm.nih.gov/blast/db on August 27, 2021. The RNAcentral database was obtained from https://ftp.ebi.ac.uk/pub/databases/RNAcentral/current_release/sequences on August 17, 2021. The MG-RAST database was established by collecting assembled transcriptomic and metagenomic sequences from https://www.mg-rast.org on October 7, 2021. The GWH database was downloaded from ftp://download.big.ac.cn/gwh on August 21, 2021. The MGnify database was downloaded from ftp://ftp.ebi.ac.uk/pub/databases/metagenomics/mgnify_genomes on December 21, 2021.

### Data processing

The NCBI databases were downloaded in NCBI-BLAST format. The corresponding fasta files were extracted by blastdbcmd from the BLAST+ 2.12.0 package[28]. The RNAcentral database is downloaded as a zipped fasta file and is used as-is after inflation. The MG-RAST, GWH, and MGnify databases are downloaded as individual sequences for assemblies. Sequences from the three sources are first merged according to their data source, resulting three bulk fasta files. The fasta files of MG-RAST and GWH are further formatted as follows: 1) sequences longer than 1000m bases (which are usually chromosomes) are deleted; 2) all sequences are transferred to DNA alphabet; 3) all gaps, dashes and non-IUPAC characters in sequences are substituted with character ‘N’. After processing, all eight databases (nt, env_nt, tsa_nt patnt, RNAcentral, MG-RAST, GWH, and MGnify) are available as eight bulk fasta files.

The above databases were concatenated in fasta format, resulting a raw total size of 1742 GB. SeqKit[29] was then employed to remove 100% duplicated sequences. The final database versioned as the MARS database 1.0. It is released in the fasta format, comprised of 1,727,789,860 nucleotide sequences with 1,592,396,862,523 bases in total and the file size reaches 1571 GB, compared to 72.9 million sequences in nt and 27 milion sequences in RNAcentral.

### Application to RNA homology search by RNAcmap

Here, we adapted the three-iteration framework of RNAcmap2 for homology search [18] with a major change how the databases were searched (Figure 1). As one large file for the sequence dataset is inefficient to handle, it was split into 149 volumes with a fixed size of 10 GB. Independent cmsearch processes in Infernal[14] are evoked on these individual volumes, producing individual multiple sequence alignments (MSAs) on the volumes. The individual MSAs are then merged into the MSA on the full database with esl-alimerge, a mini-app from Easel toolkit shipped with Infernal. This split strategy significantly improves the depth of resulting MSAs. To distinguish this change from RNAcmap2 in relation to the database search, we label the current search as RNAcmap3 against the MARS dataset for comparison with the previous RNAcmap results.

**Figure 1.**
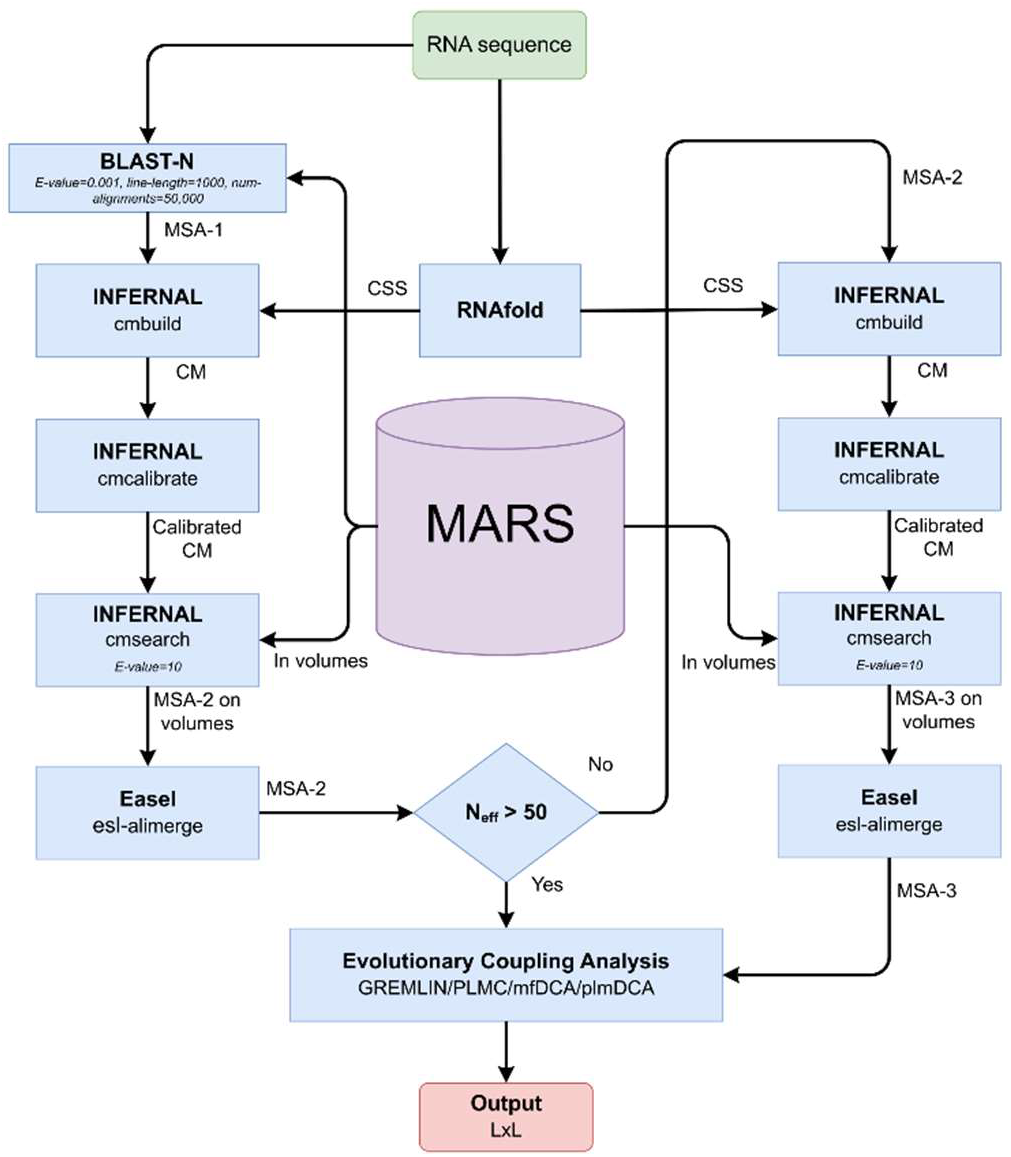
The schematic diagram of the RNAcmap3 pipeline. CSS: Consensus Secondary Structure. CM: Covariance Model. L: Length of the input RNA sequence.

### Benchmark for comparing homology searches

We employed the same benchmark datasets that were employed for comparing RNAcmap2 with RNAcmap1 [18]. Briefly, non-redundant RNA structures (80% cutoff by CD-HIT-EST [30]) were obtained from Protein Data Bank [31]. Their sequences were searched against the NCBI *nt* database by RNAcmap1 and divided into No-hit, Low N_eff_ (1-10), Medium N_eff_ (10-50) and high N_eff_ (>50) sets with 21, 83, 31, and 110 RNAs, respectively. Here, we will focus on No-hit, Low N_eff_, and Medium N_eff_ sets only because co-variational, direct coupling analysis of the MSAs for the high N_eff_ set has achieved highly accurate prediction of base pairs by RNAcmap1. More homologous sequences by RNAcmap2 or RNAcmap3 can no longer increase evolutionary or co-evolutionary information for those with high N_eff_ by RNAcmap1. The above 135 PDB structures (No-hit, Low N_eff_, and Medium N_eff_ structures) were further mapped onto Rfam and non-Rfam families by simply searching PDB RNA sequences on the Rfam website (https://rfam.xfam.org). This led to 30 different Rfam families along with 105 sequences that are not mapped to any Rfam families. The MSAs and base-pair predictions from Rfam are compared to those from RNAcmap2 as done in [18] and RNAcmap3 developed here.

The MSAs produced by RNAcmap3 were evaluated by assessing the accuracy of the secondary structure predicted by co-variational analysis of the MSAs, as in RNAcmap2 [18], according to sensitivity (SN = TP/(TP + FN)), precision (PR=TP/(TP+FP)) and F1-score (F1=2(PR ×SN/(PR+SN)) for non-local base-pairs (|i-j |>3). Here, TP, FN, and FP are true positives, false negatives, and false positives, respectively. The F1, Precision and Sensitivity are calculated with the top L/3 predictions as predicted truth. In RNAcmap2, the co-variational analysis of MSAs was done by direct coupling analysis (DCA) predictors (GREMLIN[32], mfDCA[15], PLMC[33,34] and plmDCA[35]). Because PLMC and plmDCA failed to produce the results for some MSAs generated by RNAcmap3, GREMLIN and mfDCA are utilized for method comparison. However, only mfDCA is reported here because mfDCA consistently yielded better results than GREMLIN.

rMSA is a recently reported pipeline for RNA homology search [19] that searched against nt and RNAcentral databases. Here, the versions of these two databases for rMSA were the same as used in RNAcmap3. The rMSA program is downloaded from https://github.com/pylelab/rMSA. In all searches, rMSA runs with the default run parameters.

The RNAcmap2 results presented in the manuscript are obtained on the same version of NCBI databases as used in MARS.

## RESULTS

### Performance on RNA homology search

Table 1 compares the MSAs generated by RNAcmap2, rMSA, and RNAcmap3 in term of the number of effective homologous sequences (Neff) and the F1-score given by mfDCA for the MSAs. The distribution of F1-scores for individual RNAs is shown in Figure 2.

**Table 1.**
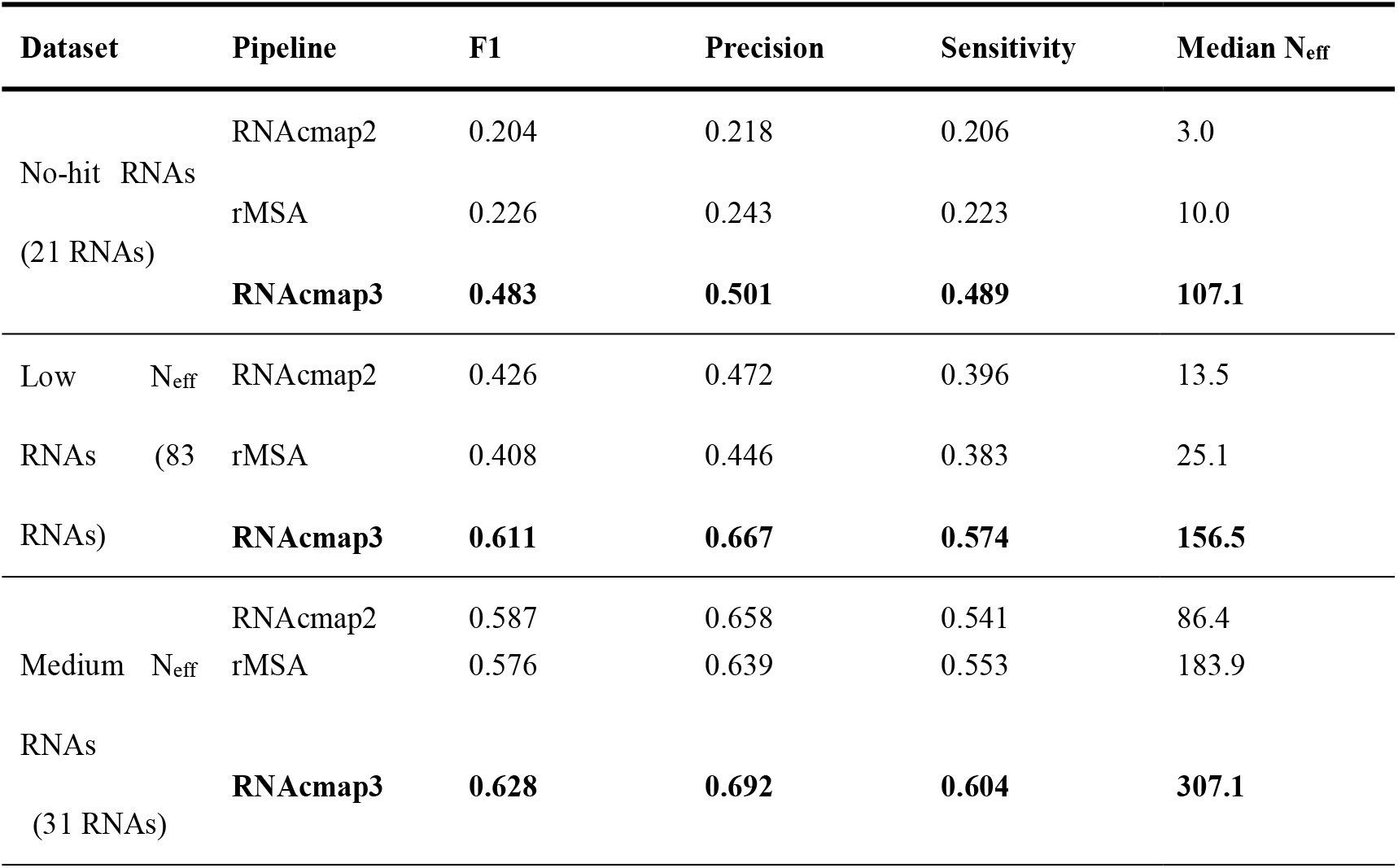
Performance comparison by F1-Score (harmonic mean of precision and sensitivity) among RNAcmap2, RNAcmap3 and rMSA on No-hit, Low N_eff_ and Medium N_eff_ datasets using mfDCA predictor.

**Figure 2.**
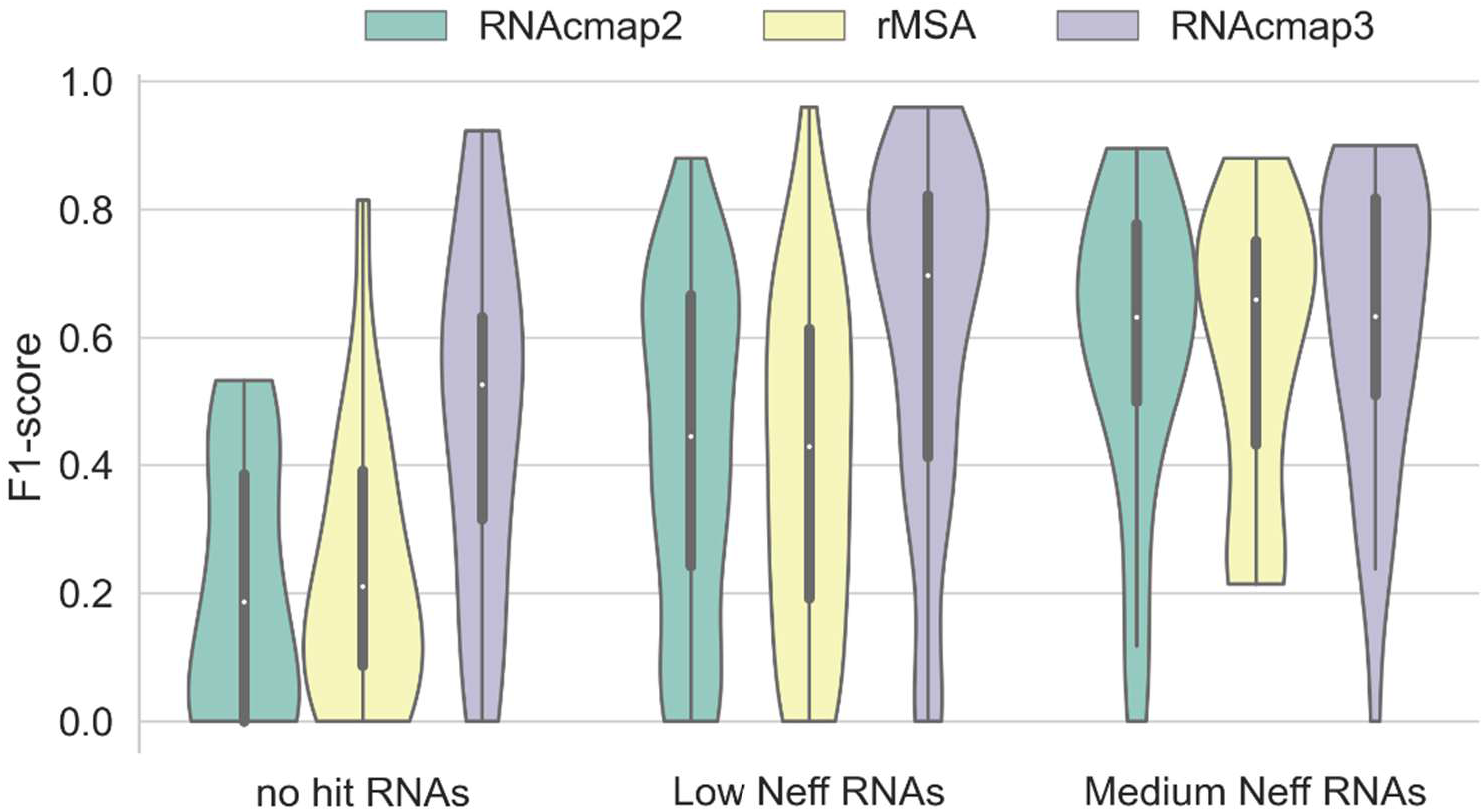
Violin plot of F1-score predicted by mfDCA using MSA generated by RNAcmap2, rMSA, and RNAcmap3. The density estimation is computed for no hit RNAs (21 RNAs), Low N_eff_ RNAs (83 RNAs) and Medium N_eff_ RNAs (31 RNAs), respectively. In the Violin plot, the empty circle denotes the median, the thick vertical bar in the centre denotes the interquartile range, and the thin vertical bar shows the range of data points within another 1.5 interquartile range extension from the thick bar ends. The Violin plot is cut off at the range of all actual data points.

RNAcmap2 and rMSA have a comparable performance in all three datasets. The average F1-score derived from rMSA results is slightly higher than RNAcmap2 in Nohit RNAs, but slightly lower in Low N_eff_ and Medium N_eff_ RNAs. This happened despite that rMSA produces MSAs with much higher average N_eff_ values than that of RNAcmap2 in all three datasets (Table 1). It seems that a higher N_eff_ value (a statistically significant difference between N_eff_ values with a p-value of 0.006) does not necessarily produce a higher MSA quality (a statistically insignificant difference between F1-scores with a p-value of 0.628).

RNAcmap3 outperforms both RNAcmap2 and rMSA in all three datasets on all performance indicators although overall comparable performance on the Medium set. RNAcmap3 increases in average F1-score over RNAcmap2 by 136.8% for no-hit RNAs, 43.4% for Low N_eff_ RNAs and 6.98% for Medium N_eff_ RNAs, respectively. RNAcmap3 also increases in average F1-score over rMSA by 113.7% for no-hit RNAs, 49.8% for Low N_eff_ RNAs and 9.0% for Medium N_eff_ RNAs, respectively. The RNAcmap3-generated MSAs have N_eff_ values much higher than that of rMSA MSAs. RNAcmap3 yields MSAs with median N_eff_ >100 even for No hit RNAs. Comparing to RNAcmap2, the high N_eff_ values (p-value=3.74 × 10^−13^) are indeed related to much better MSA qualities as reflected by the F1-scores (p-value=4.14 × 10^−7^). Note that there is a zero F1-score for RNAcmap3 (PDB 1g1x_E in Medium N_eff_ RNAs) due to poor performance of RNAfold for providing initial secondary structure employed in homology search. This leads to a smaller median F1-score for RNAcmap3 on Medium N_eff_ RNAs, compared to that for rMSA. More discussions can be found below.

Among three RNA datasets, the improvement of RNAcmap3 is the most significant for the No hit and Low N_eff_ datasets. In fact, the performance of RNAcmap3 on No hit RNAs is better than that of RNAcmap2 on Low N_eff_ RNAs. RNAcmap3 in the Low N_eff_ dataset also outperforms RNAcmap2 in the Medium N_eff_ dataset. This is consistent with the drastically increased N_eff_ values. The performance improvement of RNAcmap3 on the Medium N_eff_ RNAs is <10% over RNAcmap2 (or rMSA), because RNAcmap2 also generates MSAs with sufficient N_eff_ values. This is in line with the notion that prediction accuracy by covariational analysis along with MSA depth has an upper limit. Similar results (Supplementary Table S1 and Supplementary Figure S1) were obtained when GREMLIN was employed to measure the quality of MSA

### Comparison between RNAcmap3 and manually annotated Rfam

Rfam clusters RNA sequences into the families according to the homology in sequence and secondary structure. When possible, Rfam utilizes experimentally determined secondary structures for homology search and alignment. By comparison, a method like RNAcmap or rMSA employed RNAfold for initial secondary structure prediction. Thus, Rfam is often considered as the gold standard for RNA MSAs although not all RNAs in Rfam employed experimentally determined secondary structure.

Figure 3 shows the F1-scores from mfDCA-predicted base-pairs (top L/3) using the MSAs (1 RNA in the no-hit set, 14 in the low N_eff_ set and 15 in the medium N_eff_ set) from Rfam, RNAcmap2, and RNAcmap3, respectively. For medium N_eff_ RNAs, RNAcmap3 retains the significantly improved performance of RNAcmap2 over Rfam. For low N_eff_ RNAs, the performance on some sequences is significantly improved over Rfam (and RNAcmap2) but not improved for others.

**Figure 3.**
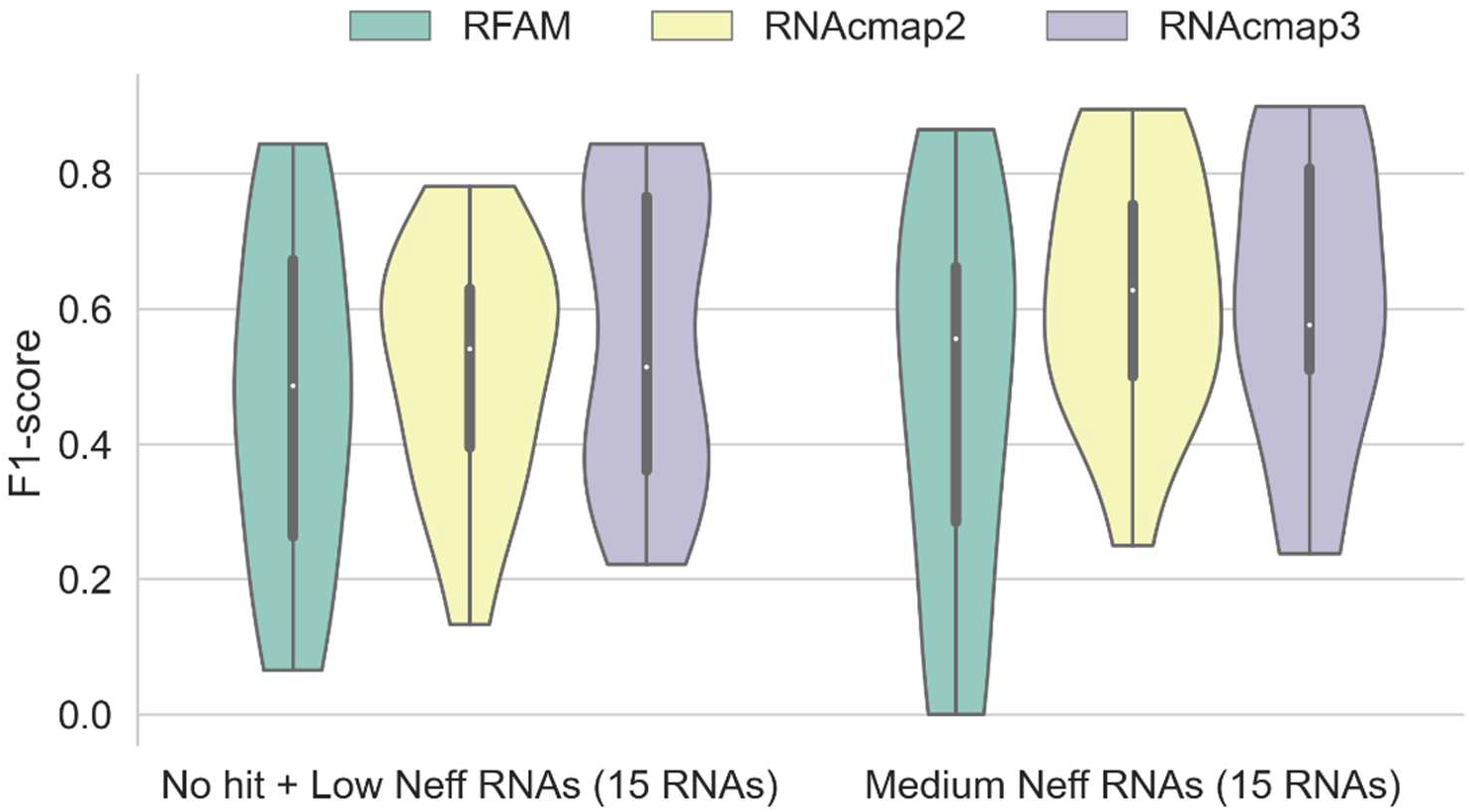
Violin plot of F1-score predicted by mfDCA using MSAs provided by Rfam, RNAcmap2 and RNAcmap3 for RNAs mapped to Rfam. The density estimation is computed for no hit RNAs (1 RNAs), Low Neff RNAs (14 RNAs), and Medium Neff RNAs (15 RNAs), respectively. In the Violin plot, the empty circle denotes the median, the thick vertical bar in the centre denotes the interquartile range, and the thin vertical bar shows the range of data points within another 1.5 interquartile range extension from the thick bar ends. The Violin plot is cut off at the range of all actual data points.

A more detailed comparison for each family is shown in Table 2. In the mapped 30 families, RNAcmap3 outperforms Rfam in 17 families, and Rfam performs better than RNAcmap3 in 10 families, with equal performance on 3 families. On the other hand, RNAcmap3 outperforms RNAcmap2 in 15/30 families and RNAcmap2 outperforms RNAcmap3 in 9/30 families, with 6 families in similar performance. RNAcmap3 improves more over RNAcmap2 when both improves over Rfam (10 in 18 families). In the 9 families that RNAcmap2 does not perform as well as Rfam, RNAcmap3 improves the performance in 5 families, while fails to do so on the remaining 4. Interestingly, average-speaking, RNAcmap2 performs better on the Rfam-mapped sequences than Non-Rfam sequences. However, RNAcmap3 seems to perform even better on Non-Rfam sequences than the Rfam-mapped sequences.

**Table 2.**
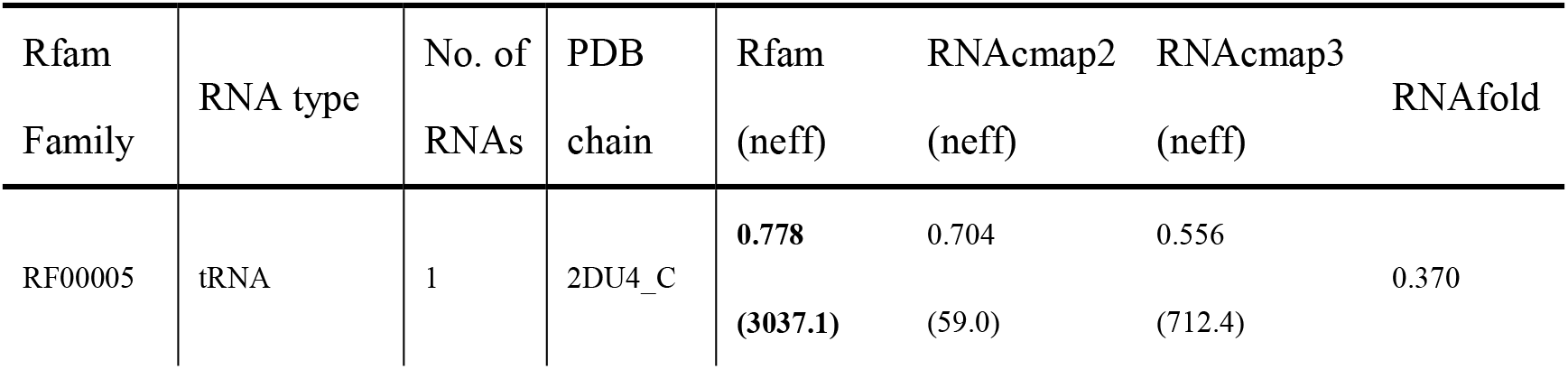
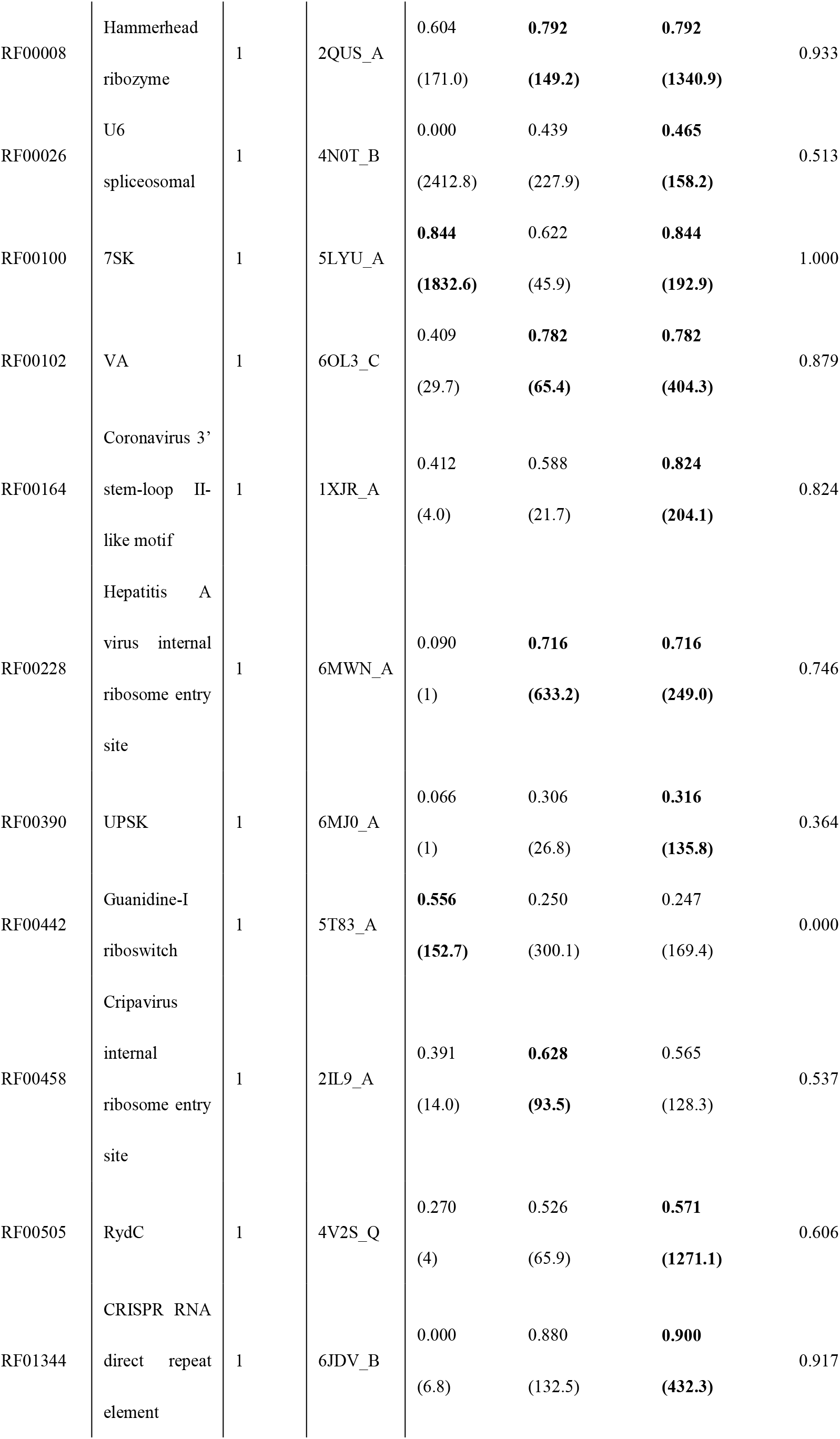
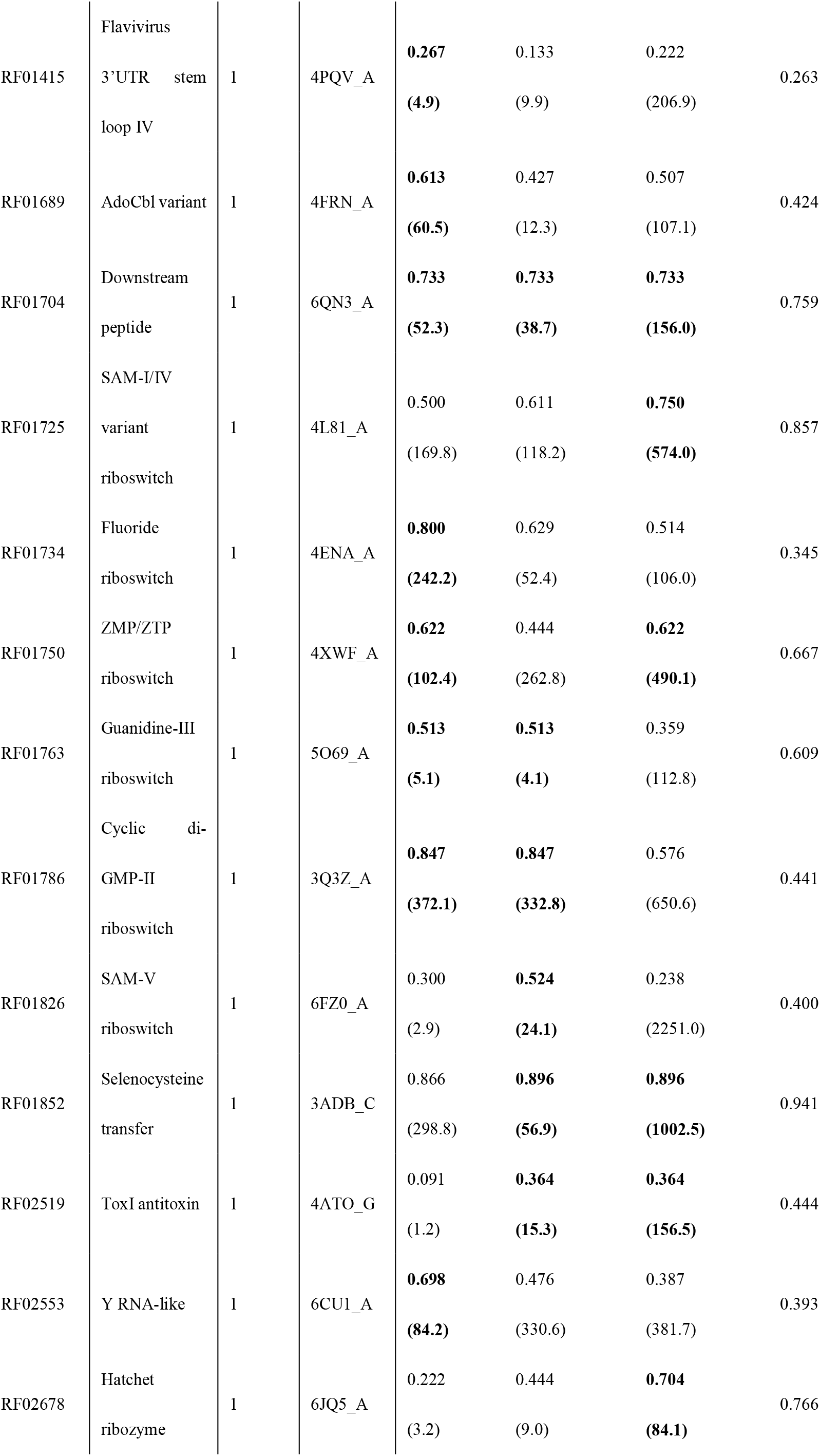
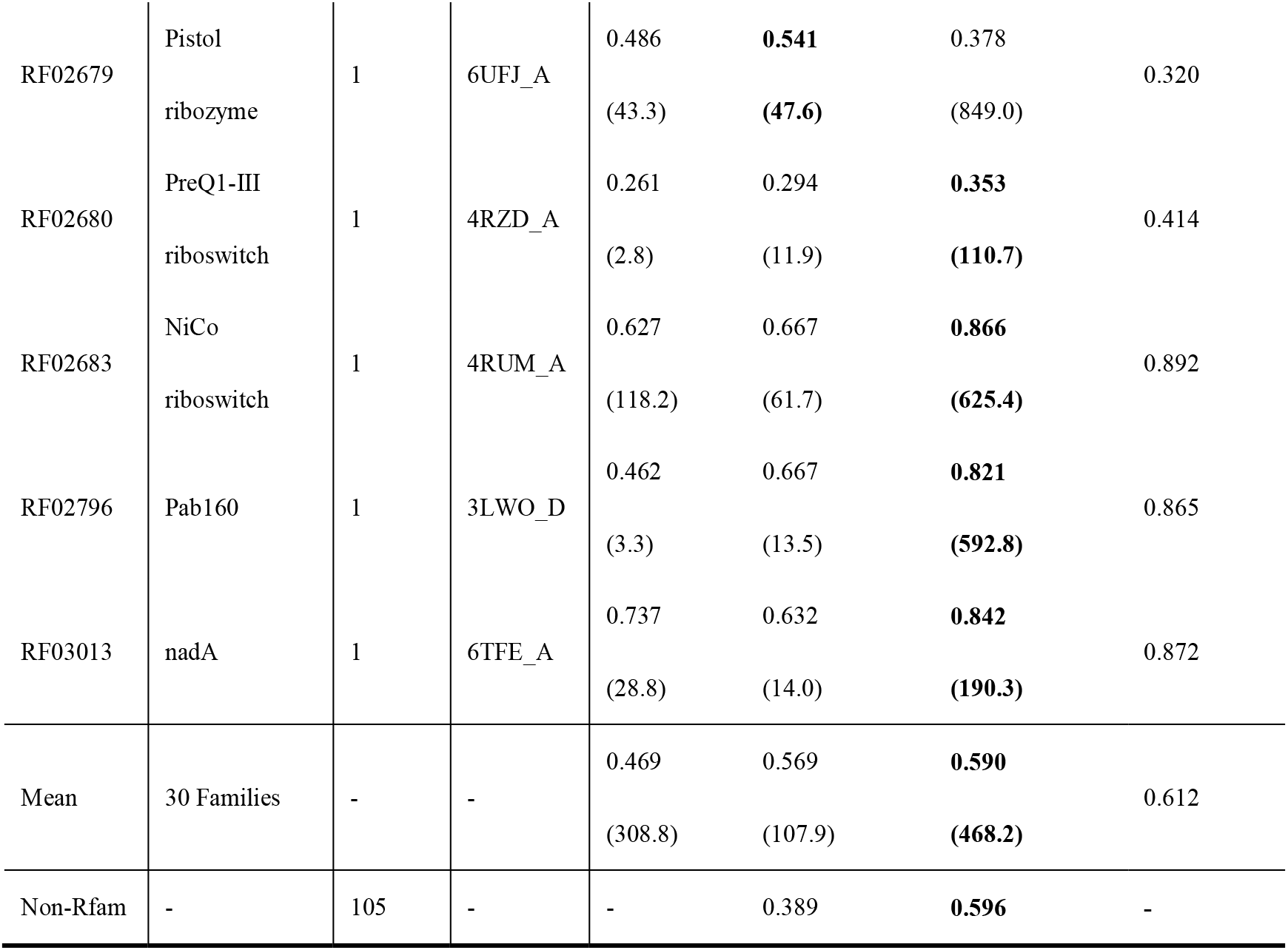
Performance given by Rfam, RNAcmap2, RNAcmap3 and RNAfold with F1-score on 30 Rfam mapped families using the mfDCA predictor, as well as on 105 non-Rfam RNAs.

One big difference between Rfam and RNAcmap is that Rfam relied on known secondary structures whereas RNAcmap employed secondary structure predicted by RNAfold. Table 2 and Figure 4 illustrated the dependence of RNAcmap performance on RNAfold. In particular, the improvement of RNAcmap2 or RNAcmap3 over Rfam F1 score is positively correlated with the F1 score given by RNAfold with a Pearson’s correlation coefficient of 0.573 (p=0.001) for RNAcmap3 and 0.359 (p=0.051) for RNAcmap2. If RNAfold predictions have a F1-score of >0.667, RNAcmap3 always performs equally or better than RNAcmap2 and Rfam.

**Figure 4.**
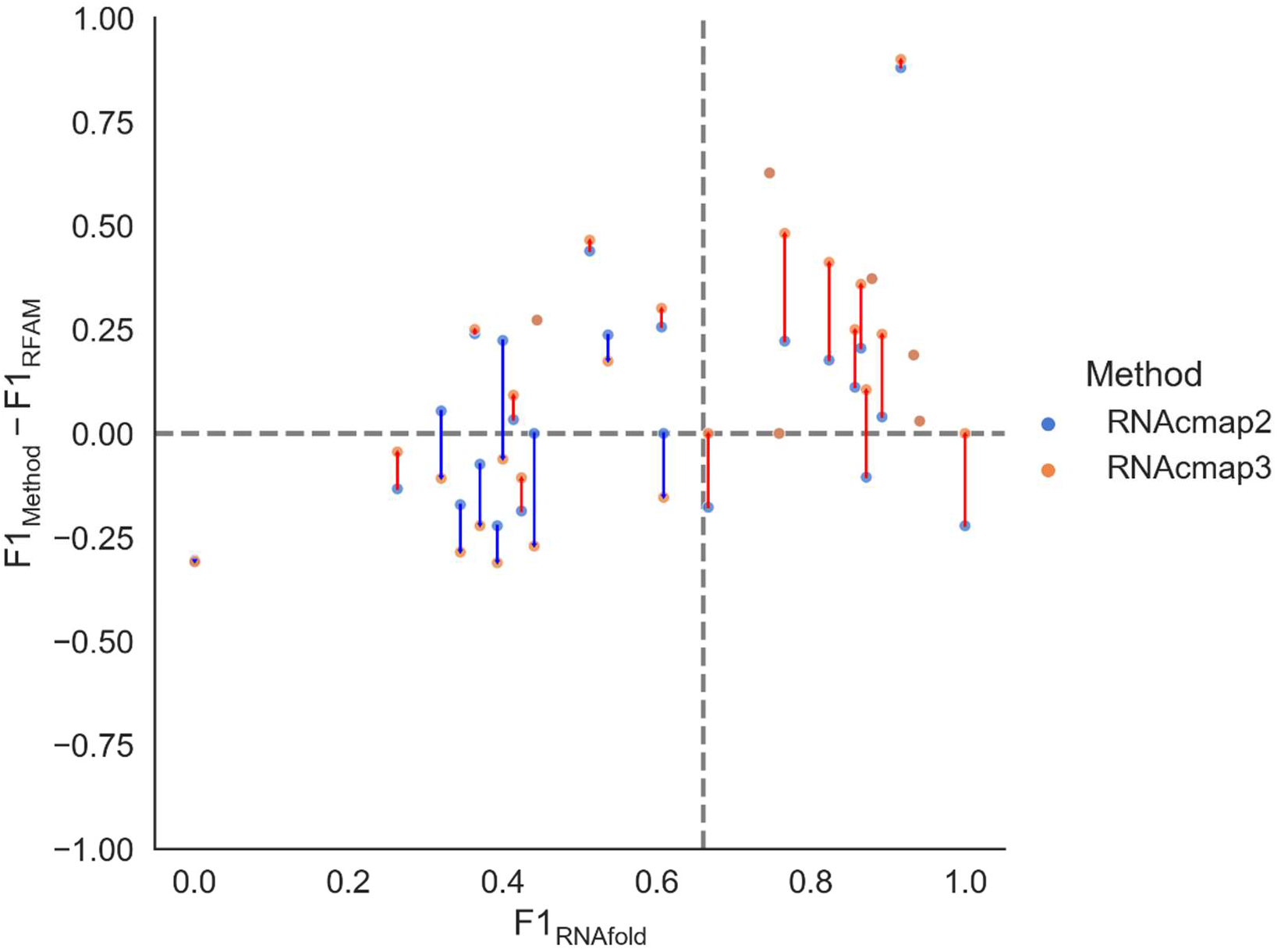
The difference between F1-scores given by RNAcmap2 (or RNAcmap3) and the F1-scores given by Rfam as a function of F1-score given by RNAfold. RNAcmap3 and RNAcmap2 results are shown in orange and blue, respectively. The results of RNAcmap3 and RNAcmap2 are linked with a red line if RNAcmap3 improves over RNAcmap2 and a blue line, if otherwise.

## DISCUSSION

This work established a comprehensive database of nucleotide sequences by including NCBI’s nucleotide database (nt), environmental samples (env_nt), transcriptome shotgun assembly (tsa_nt) and nucleotide sequences from the Patent Division of Genbank (patnt), the noncoding RNA sequences from RNAcentral, the transcriptome assembly and metagenome assembly from MG-RAST, the genomic sequences from Genome Warehouse (GWH) and the genomic sequences from MGnify. This compilation led to the MARS database of nucleotide sequences that is more than 20 times larger than the commonly used nt database in the number of sequences. Using a split-search strategy for the MARS database allows RNAcmap3 to gain a deeper MSA and yield better co-evolution coupling than RNAcmap2 and rMSA. Moreover, despite using RNAfold as the initial secondary structure for homology inference, RNAcmap3 can achieve more accurate inference of secondary structure from MSA than from Rfam MSAs. RNAcmap3 is expected to be useful for improving RNA homology search.

One issue of MARS is the huge size of the sequence datasets with 1.5TB for its first version. This huge size makes the homology search very slow, despite of the strategy of data splitting for parallel processing. A typical search for a 100-base long sequence would take 4 hours on 24 cpus. Longer sequences of >1000 nucleotides long are prohibitively slow. One expects that the sequence database will continue to expand exponentially given the low cost of high-throughput sequencing. Unfortunately, not all datasets contained in the MARS can be updated fully automatically. For example, an ftp access to the MGnify database with a script frequently suffers from broken connections. One must rely on manual intervention to complete the process.

For RNAcmap3, one limitation is that one must use a predicted secondary structure as the initial guess for homology search. Here, we employed RNAfold. We found that the performance of the method is somewhat depending on how accurate is the initial RNAfold prediction (Figure 4). This problem can be addressed with improved prediction of secondary structure, for example, by deep learning techniques (e.g. SPOT-RNA [36], MXfold2[37], UFold [38]). However, there is a risk of overtraining for some of these deep learning techniques, which would make some methods to perform poorly for unseen RNA families [39]. Thus, caution must be exercised when using these deep learning techniques.

## Data availability

All datasets, RNAcmap3 can be downloaded from http://zhouyq-lab.szbl.ac.cn/download/

## Acknowledgement

The authors gratefully acknowledge that the High Performance Computing Cluster at Shenzhen Bay Laboratory was involved in completing this research. The support of National Key Research and Development Program of China(NO.2021YFF1200400) and the Major Program of Shenzhen Bay Laboratory S201101001, and Shenzhen Science and Technology Program [KQTD20170330155106581] are acknowledged.

## Supplementary material

**Table S1 Performance comparison between RNAcmap2, RNAcmap3 and rMSA on No-hit, Low Neff and Medium Neff datasets using the GREMLIN predictor**

**Figure S1 Violin plot of F1-score predicted by GREMLIN using MSA generated by RNAcmap2, rMSA and RNAcmap3.** The density estimation is computed for no hit RNAs (21 RNAs), Low N_eff_ RNAs (83 RNAs), and Medium N_eff_ RNAs (31 RNAs), respectively. In the Violin plot, the empty circle denotes the median, the thick vertical bar in the centre denotes the interquartile range, and the thin vertical bar shows the range of data points that within another 1.5 interquartile range extension from the thick bar ends. The Violin plot is cut off at the range of all actual data points.

## References

[1] Hoagland MB, Stephenson ML, Scott JF, Hecht LI, Zamecnik PC. A soluble ribonucleic acid intermediate in protein synthesis. J Biol Chem 1958;231:241–57.

[2] Fabbri M, Girnita L, Varani G, Calin GA. Decrypting noncoding RNA interactions, structures, and functional networks. Genome Res 2019;29:1377–88. https://doi.org/10.1101/gr.247239.118.

[3] Bushati N, Cohen SM. microRNA functions. Annu Rev Cell Dev Biol 2007;23:175–205. https://doi.org/10.1146/annurev.cellbio.23.090506.123406.

[4] Sleutels F, Zwart R, Barlow DP. The non-coding Air RNA is required for silencing autosomal imprinted genes. Nature 2002;415:810–3. https://doi.org/10.1038/415810a.

[5] Micura R, Höbartner C. On secondary structure rearrangements and equilibria of small RNAs. Chembiochem 2003;4:984–90. https://doi.org/10.1002/cbic.200300664.

[6] Westhof E, Leontis NB. An RNA-centric historical narrative around the Protein Data Bank. J Biol Chem 2021;296:100555. https://doi.org/10.1016/j.jbc.2021.100555.

[7] Bertone P, Stolc V, Royce TE, Rozowsky JS, Urban AE, Zhu X, et al. Global identification of human transcribed sequences with genome tiling arrays. Science 2004;306:2242–6. https://doi.org/10.1126/science.1103388.

[8] Zhou B, Ji B, Liu K, Hu G, Wang F, Chen Q, et al. EVLncRNAs 2.0: an updated database of manually curated functional long non-coding RNAs validated by low-throughput experiments. Nucleic Acids Res 2021;49:D86–91. https://doi.org/10.1093/nar/gkaa1076.

[9] RNAcentral Consortium. RNAcentral 2021: secondary structure integration, improved sequence search and new member databases. Nucleic Acids Research 2021;49:D212–20. https://doi.org/10.1093/nar/gkaa921.

[10] Sayers EW, Beck J, Bolton EE, Bourexis D, Brister JR, Canese K, et al. Database resources of the National Center for Biotechnology Information. Nucleic Acids Res 2021;49:D10–7. https://doi.org/10.1093/nar/gkaa892.

[11] Jumper J, Evans R, Pritzel A, Green T, Figurnov M, Ronneberger O, et al. Highly accurate protein structure prediction with AlphaFold. Nature 2021. https://doi.org/10.1038/s41586-021-03819-2.

[12] Zhang T, Singh J, Litfin T, Zhan J, Paliwal K, Zhou Y. RNAcmap: a fully automatic pipeline for predicting contact maps of RNAs by evolutionary coupling analysis. Bioinformatics 2021. https://doi.org/10.1093/bioinformatics/btab391.

[13] Altschul SF, Madden TL, Schäffer AA, Zhang J, Zhang Z, Miller W, et al. Gapped BLAST and PSI-BLAST: a new generation of protein database search programs. Nucleic Acids Research 1997;25:3389–402. https://doi.org/10.1093/nar/25.17.3389.

[14] Nawrocki EP, Eddy SR. Infernal 1.1: 100-fold faster RNA homology searches. Bioinformatics 2013;29:2933–5. https://doi.org/10.1093/bioinformatics/btt509.

[15] Morcos F, Pagnani A, Lunt B, Bertolino A, Marks DS, Sander C, et al. Direct-coupling analysis of residue coevolution captures native contacts across many protein families. Proceedings of the National Academy of Sciences 2011;108:E1293–301. https://doi.org/10.1073/pnas.1111471108.

[16] Singh J, Paliwal K, Zhang T, Singh J, Litfin T, Zhou Y. Improved RNA secondary structure and tertiary base-pairing prediction using evolutionary profile, mutational coupling and two-dimensional transfer learning. Bioinformatics 2021;37:2589–600. https://doi.org/10.1093/bioinformatics/btab165.

[17] Singh J, Paliwal K, Litfin T, Singh J, Zhou Y. Predicting RNA distance-based contact maps by integrated deep learning on physics-inferred secondary structure and evolutionary-derived mutational coupling. Bioinformatics 2022:btac421. https://doi.org/10.1093/bioinformatics/btac421.

[18] Singh J, Paliwal K, Singh J, Litfin T, Zhou Y. Improved RNA homology detection and alignment by automatic iterative search in an expanded database 2022:2022.10.03.510702. https://doi.org/10.1101/2022.10.03.510702.

[19] Zhang C, Zhang Y, Pyle AM. rMSA: A Sequence Search and Alignment Algorithm to Improve RNA Structure Modeling. Journal of Molecular Biology 2022:167904. https://doi.org/10.1016/j.jmb.2022.167904.

[20] Pearce R, Omenn GS, Zhang Y. De Novo RNA Tertiary Structure Prediction at Atomic Resolution Using Geometric Potentials from Deep Learning 2022:2022.05.15.491755. https://doi.org/10.1101/2022.05.15.491755.

[21] Kalvari I, Nawrocki EP, Ontiveros-Palacios N, Argasinska J, Lamkiewicz K, Marz M, et al. Rfam 14: expanded coverage of metagenomic, viral and microRNA families. Nucleic Acids Research 2021;49:D192–200. https://doi.org/10.1093/nar/gkaa1047.

[22] Wheeler TJ, Eddy SR. nhmmer: DNA homology search with profile HMMs. Bioinformatics 2013;29:2487–9. https://doi.org/10.1093/bioinformatics/btt403.

[23] Meyer F, Paarmann D, D’Souza M, Olson R, Glass E, Kubal M, et al. The metagenomics RAST server – a public resource for the automatic phylogenetic and functional analysis of metagenomes. BMC Bioinformatics 2008;9:386. https://doi.org/10.1186/1471-2105-9-386.

[24] Wilke A, Bischof J, Harrison T, Brettin T, D’Souza M, Gerlach W, et al. A RESTful API for Accessing Microbial Community Data for MG-RAST. PLOS Computational Biology 2015;11:e1004008. https://doi.org/10.1371/journal.pcbi.1004008.

[25] Chen M, Ma Y, Wu S, Zheng X, Kang H, Sang J, et al. Genome Warehouse: A Public Repository Housing Genome-scale Data. Genomics, Proteomics & Bioinformatics 2021;19:584–9. https://doi.org/10.1016/j.gpb.2021.04.001.

[26] CNCB-NGDC Members and Partners. Database Resources of the National Genomics Data Center, China National Center for Bioinformation in 2022. Nucleic Acids Research 2022;50:D27–38. https://doi.org/10.1093/nar/gkab951.

[27] Mitchell AL, Almeida A, Beracochea M, Boland M, Burgin J, Cochrane G, et al. MGnify: the microbiome analysis resource in 2020. Nucleic Acids Research 2020;48:D570–8. https://doi.org/10.1093/nar/gkz1035.

[28] Camacho C, Coulouris G, Avagyan V, Ma N, Papadopoulos J, Bealer K, et al. BLAST+: architecture and applications. BMC Bioinformatics 2009;10:421. https://doi.org/10.1186/1471-2105-10-421.

[29] Shen W, Le S, Li Y, Hu F. SeqKit: A Cross-Platform and Ultrafast Toolkit for FASTA/Q File Manipulation. PLOS ONE 2016;11:e0163962. https://doi.org/10.1371/journal.pone.0163962.

[30] Li W, Godzik A. Cd-hit: a fast program for clustering and comparing large sets of protein or nucleotide sequences. Bioinformatics 2006;22:1658–9. https://doi.org/10.1093/bioinformatics/btl158.

[31] Burley SK, Bhikadiya C, Bi C, Bittrich S, Chen L, Crichlow GV, et al. RCSB Protein Data Bank: powerful new tools for exploring 3D structures of biological macromolecules for basic and applied research and education in fundamental biology, biomedicine, biotechnology, bioengineering and energy sciences. Nucleic Acids Research 2021;49:D437–51. https://doi.org/10.1093/nar/gkaa1038.

[32] Kamisetty H, Ovchinnikov S, Baker D. Assessing the utility of coevolution-based residue–residue contact predictions in a sequence-and structure-rich era. Proceedings of the National Academy of Sciences 2013;110:15674–9. https://doi.org/10.1073/pnas.1314045110.

[33] Balakrishnan S, Kamisetty H, Carbonell JG, Lee S-I, Langmead CJ. Learning generative models for protein fold families. Proteins: Structure, Function, and Bioinformatics 2011;79:1061–78. https://doi.org/10.1002/prot.22934.

[34] Hopf TA, Ingraham JB, Poelwijk FJ, Schärfe CPI, Springer M, Sander C, et al. Mutation effects predicted from sequence co-variation. Nat Biotechnol 2017;35:128–35. https://doi.org/10.1038/nbt.3769.

[35] Ekeberg M, Lövkvist C, Lan Y, Weigt M, Aurell E. Improved contact prediction in proteins: Using pseudolikelihoods to infer Potts models. Phys Rev E 2013;87:012707. https://doi.org/10.1103/PhysRevE.87.012707.

[36] Singh J, Hanson J, Paliwal K, Zhou Y. RNA secondary structure prediction using an ensemble of two-dimensional deep neural networks and transfer learning. Nat Commun 2019;10:5407. https://doi.org/10.1038/s41467-019-13395-9.

[37] Sato K, Akiyama M, Sakakibara Y. RNA secondary structure prediction using deep learning with thermodynamic integration. Nat Commun 2021;12:941. https://doi.org/10.1038/s41467-021-21194-4.

[38] Fu L, Cao Y, Wu J, Peng Q, Nie Q, Xie X. UFold: fast and accurate RNA secondary structure prediction with deep learning. Nucleic Acids Res 2022;50:e14. https://doi.org/10.1093/nar/gkab1074.

[39] Szikszai M, Wise M, Datta A, Ward M, Mathews DH. Deep learning models for RNA secondary structure prediction (probably) do not generalize across families. Bioinformatics 2022;38:3892–9. https://doi.org/10.1093/bioinformatics/btac415.

