## Supplementary Materials for "The Master Database of All Possible RNA Sequences and Its Integration with RNAcmap for RNA Homology Search"

[†]Co-first authors.

**Table S1 Performance comparison between RNAcmap2, RNAcmap3, and rMSA on No-hit, Low Neff, and Medium Neff datasets using the GREMLIN predictor**

| Dataset | Pipeline | F1 | Precision | Sensitivity | Median $N_{eff}$ |
|---|---|---|---|---|---|
| No-hit RNAs (21 RNAs) | RNAcmap2 | 0.187 | 0.198 | 0.195 | 3.0 |
| | RNAcmap3 | **0.466** | **0.484** | **0.469** | **107.1** |
| | rMSA | 0.177 | 0.193 | 0.169 | 10.0 |
| Low $N_{eff}$ RNAs (83 RNAs) | RNAcmap2 | 0.368 | 0.405 | 0.347 | 13.5 |
| | RNAcmap3 | **0.584** | **0.640** | **0.546** | **156.5** |
| | rMSA | 0.381 | 0.417 | 0.357 | 25.1 |
| Medium $N_{eff}$ RNAs (31 RNAs) | RNAcmap2 | 0.518 | 0.583 | 0.476 | 86.4 |
| | RNAcmap3 | **0.584** | **0.646** | **0.560** | **307.1** |
| | rMSA | 0.490 | 0.547 | 0.471 | 183.9 |

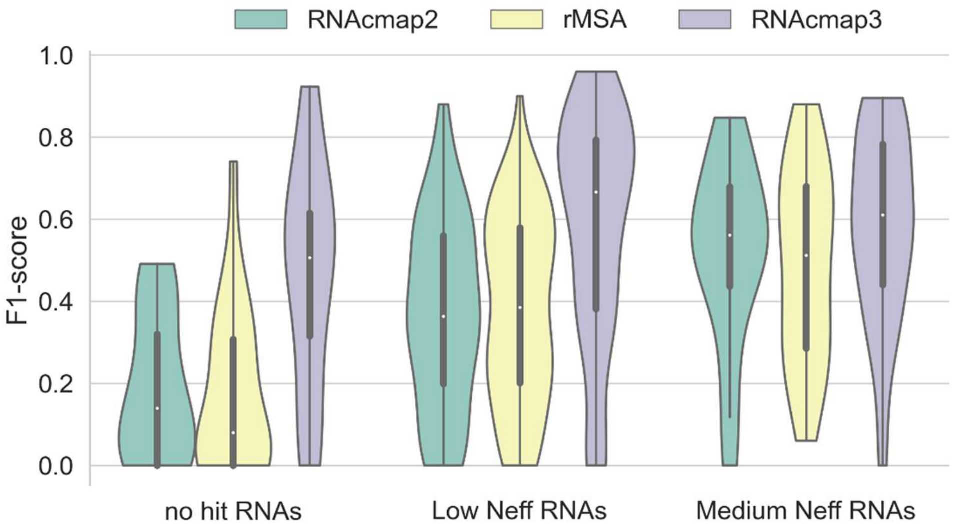


**Figure S1 Violin plot of F1-score predicted by GREMLIN using MSA generated by RNAcmap2, rMSA, and RNAcmap3.** The density estimation is computed for no hit RNAs (21 RNAs), Low $N_{eff}$ RNAs (83 RNAs), and Medium $N_{eff}$ RNAs (31 RNAs), respectively. In the Violin plot, the empty circle denotes the median, the thick vertical bar in the centre denotes the interquartile range, and the thin vertical bar shows the range of data points within another 1.5 interquartile range extension from the thick bar ends. The Violin plot is cut off at the range of all actual data points.
